# Rational design of a new class of protease inhibitors for the potential treatment of coronavirus diseases

**DOI:** 10.1101/2020.09.15.275891

**Authors:** Michael Westberg, Yichi Su, Xinzhi Zou, Lin Ning, Brett Hurst, Bart Tarbet, Michael Z. Lin

**Affiliations:** Department of Neurobiology, Stanford University, Stanford, CA 94305, USA; Department of Chemistry, Aarhus University, Aarhus, Denmark; Department of Bioengineering, Stanford University, Stanford, CA 94305, USA; Institute for Antiviral Research, Utah State University, Logan, UT 84322, USA; Department of Animal, Dairy, and Veterinary Sciences, Utah State University, Logan, UT 84322, USA; Department of Chemical and Systems Biology, Stanford University, Stanford, CA 94305, USA

## Abstract

The coronavirus main protease, M^pro^, is a key protein in the virus life cycle and a major drug target. Based on crystal structures of SARSCoV2 M^pro^ complexed with peptidomimetic inhibitors, we recognized a binding characteristic shared with proline-containing inhibitors of hepatitis C virus protease. Initial tests showed that this subclass of HCV protease inhibitors indeed exhibited activity against M^pro^. Postulating a benefit for a preorganized backbone conformation, we designed new ketoamide-based M^pro^ inhibitors based on central proline rings. One of the designed compounds, ML1000, inhibits M^pro^ with low-nanomolar affinity and suppresses SARSCoV2 viral replication in human cells at sub-micromolar concentrations. Our findings identify ML1000 as a promising new pre-organized scaffold for the development of anti-coronavirus drugs.

## INTRODUCTION

In the last two decades, three coronaviruses, SARSCoV1, MERSCoV, and SARSCoV2 have emerged from animal reservoirs to cause lethal respiratory illnesses in humans (*1*). In particular, SARSCoV2 has already caused more than 800,000 deaths (*2*). For SARSCoV2 and future coronavirus outbreaks, until effective vaccines have been administered to most of the world’s population, antiviral drugs will be needed to reduce morbidity and death.

Orally available anti-coronavirus drugs would be highly desirable, as they can be given in an outpatient setting, i.e. without requiring hospitalization. As such they can be given upon diagnosis, which could prevent progression to more severe or protracted disease and thereby greatly reduce morbidity and mortality. However, the only antiviral drug with proven efficacy against SARSCoV2, remdesivir, is an intravenous drug that can only be administered to hospitalized patients. Even then it only reduces disease duration and risk of death by ∼30% each (*3*). Dexamethasone has proven efficacy in improving survival in late-stage COVID-19 by suppressing the late hyperinflammatory responses that cause respiratory and organ failure rather than inhibiting viral replication (*4*). However, it is ineffective if given earlier (*4*), and even possibly harmful by inhibiting immune responses directed against the virus (*5–7*). Thus, drugs that can effectively suppress coronavirus replication in an outpatient setting during the entire post-infection period do not exist, but are urgently needed.

Inhibitors of the coronavirus main protease (M^pro^), a key protein in the coronavirus life cycle, could be especially effective as antiviral agents (*8, 9*). M^pro^ (also called 3C-like protease due to its homology with enterovirus 3C protease) is a cysteine protease that cleaves the large nonstructural polyprotein of coronavirus into smaller pieces to assemble the RNA replicase (*10*). Experimental inhibitors of coronavirus M^pro^ protect mice and cats from lethal coronavirus disease, validating M^pro^ as a medicinal target (*11–13*).

There is currently intense interest in discovering effective inhibitors for SARSCoV2 M^pro^. As coronavirus M^pro^ species all share similar substrate specificity profiles (*14*), previously characterized inhibitors for other coronaviruses have, unsurprisingly, demonstrated inhibitory activity on SARSCoV2 M^pro^ as well. For example, the Michael acceptor compound N3, developed in 2005 as an inhibitor of SARSCoV1 M^pro^ (*15*), also inhibits the 96% identical SARSCoV2 M^pro^ *in vitro* (*15, 16, 16*). A series of ketoamide-based inhibitors developed for SARSCoV1 M^pro^ were also effective against SARSCoV2 M^pro^ *in vitro* (*17*), and gave rise to the modified SARSCoV2 M^pro^ inhibitor 13b (*18*). Finally, GC376, a broad-spectrum aldehyde prodrug of various coronavirus M^pro^ species, and 11a, an aldehyde compound designed to inhibit SARSCoV1 M^pro^ were found to inhibit SARSCoV2 M^pro^ (*19, 20*).

However, it is not clear if known coronavirus M^pro^ inhibitors have the necessary attributes to be safe and effective oral therapeutics for human coronavirus diseases. In 11a and the active form of GC-376, an aldehyde reactive group (warhead) reacts with the active site cysteine, but aldehydes are generally considered non-ideal drug candidates due to toxicity caused by off-target reactions (*21–24*). For example, the high reactivity of aldehyde warheads has been shown to drive non-specific inhibition of cysteine proteases (*25*). Protease inhibitors with ketoamide and Michael acceptor warheads can exhibit better specificity, stability, and oral bioavailability than those with aldehydes (*26–29*), and examples have progressed to late-stage clinical trials or regulatory approval (*30, 31*). Compounds 13b and N3 utilize ketoamide and Michael acceptor warheads respectively, but they appear to have insufficient efficacy to be therapeutically useful. While 13b demonstrates a 50% inhibitory concentration (IC_50_) on M^pro^ *in vitro* of 670 nM, the 50% effective concentration (EC_50_) in blocking SARSCoV2 replication in human cells is ∼4 μM (*18*). This compares unfavorably to the achieved concentration in mouse lungs of <0.1 μM (*18*). For N3, while it demonstrates a low IC_50_ *in vitro*, its EC_50_ for SARSCoV2 replication is 17 μM (*16*), indicating poor cell permeability. This is not surprising given its large size (mw 681) and numerous polar groups. Thus, the development of new M^pro^ inhibitor designs with low toxicity and low EC_50_ in cells remains an urgent priority.

Multiple clinically approved protease inhibitors incorporate cyclic structures to improve affinity by pre-organizing the inhibitors into a conformation favorable for binding (*32*). In energetic terms, this is considered paying the entropic penalty of conformational rigidification prior to binding (*33, 34*), and cyclization has been experimentally confirmed to increase protease inhibitor affinity (*35*). In this study, we explore the ability of a cyclic moiety to improve the affinity of coronavirus M^pro^ inhibitors. We developed a novel inhibitor, ML1000, with molecular weight of 549 Da and IC_50_ value of 12 nM on SARSCoV2 M^pro^. ML1000 inhibits SARSCoV2 replication in human cells with an EC_50_ of 0.1 μM, and thus is the highest-potency non-aldehyde M^pro^ inhibitor reported so far. ML1000 represents a new structural class of potent ketoamide inhibitors of coronavirus M^pro^, and may serve as a promising starting point for the development of drug candidates against human coronavirus diseases.

## RESULTS

While visualizing the co-crystal structure of 13b and SARSCoV2 M^pro^ (*18*), we noticed that 13b adopts a pronounced kink in its main chain at the P2 residue, i.e. the 2^nd^ residue N-terminal to the scissile bond facing the enzyme’s S2 pocket **(Fig. 1A)**. Due to our previous work with HCV protease and its small-molecule inhibitors (*36*), we recognized this kink to be similar to that created by proline analog rings in the clinically approved anti-HCV drugs boceprevir, narlaprevir, and telaprevir (**Fig. 1B-D**). Like 13b, these drugs are ketoamide-based covalent inhibitors, serving as substrates for nucleophilic attack by the deprotonated active-site cysteine.

**Figure 1.**
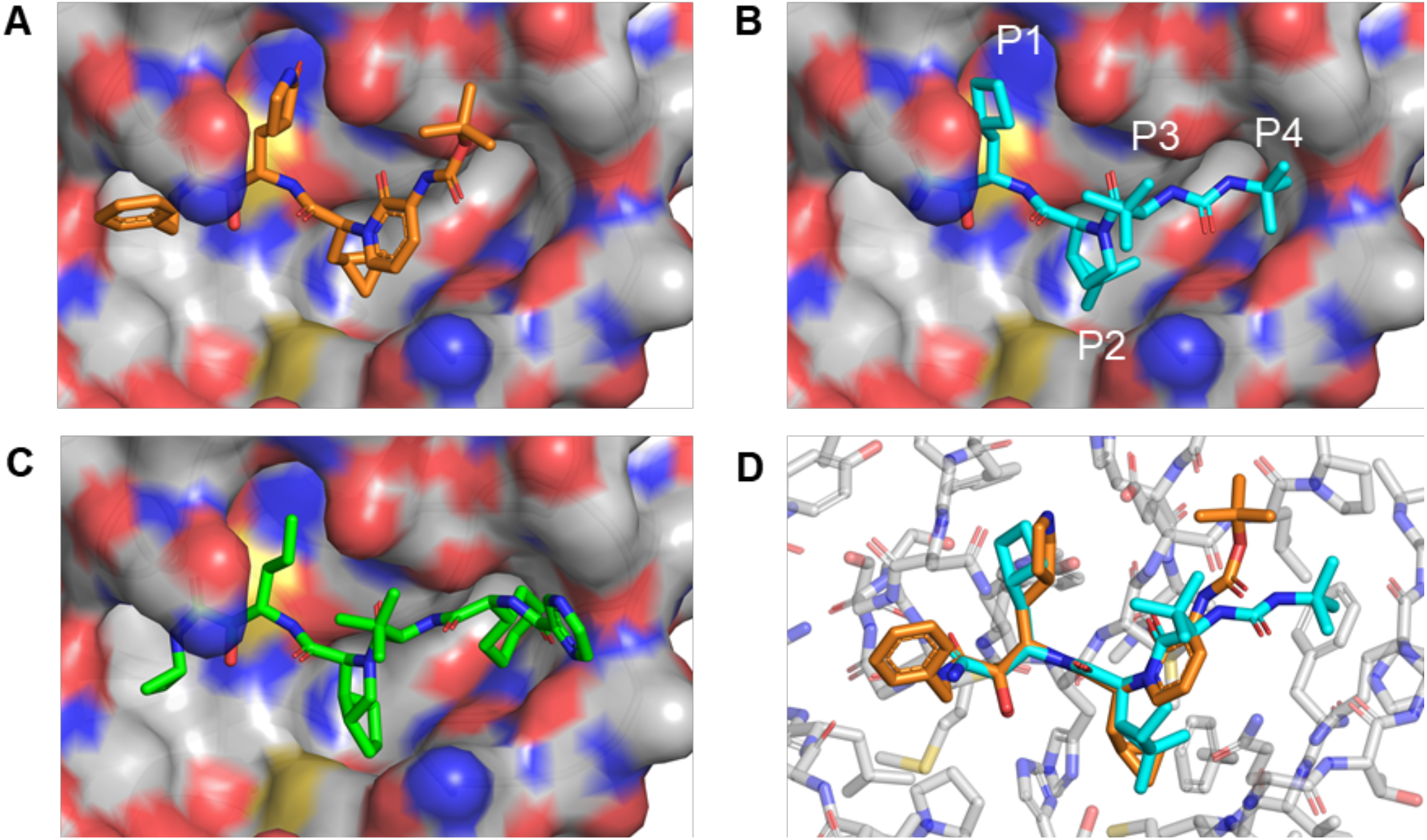
HCV protease inhibitors with a P2 proline analog can be docked into the SARSCoV2 M^pro^ active site. **(A)** Co-crystal structure of SARSCoV2 M^pro^ and inhibitor 13b (PDB 6Y2G). **(B)** Using Pymol, boceprevir was placed into the SARSCoV2 M^pro^ active site and unconstrained bonds were manually rotated for optimal complementary with the S1 and S2 pockets and hydrogen-bonding to the backbone carbonyl of Glu-166. **(C)** Telaprevir was similarly docked into the SARSCoV2 M^pro^ active site for optimal complementary with the S1, S2, and S4 pockets and hydrogen-bonding to the backbone carbonyl of Glu-166. **(D)** Alignment of the 13b-M^pro^ cocrystal with the manually docked boceprevir structure shows that the backbone of the P2-analogous segment of 13b is superimposable with the proline analog of boceprevir.

Manual rigid docking of boceprevir showed that it could fit into the active site of SARSCoV2 M^pro^ with good shape complementarity by its P1, P2, and P4 groups (**Fig. 1B**). In addition, the urea group of boceprevir at the P3-P4 junction could engage in a bidentate hydrogen bond with the backbone carbonyl of M^pro^ Glu-166. The boceprevir derivative narlaprevir also demonstrated complementary in its P1 and P2 groups, which are identical to boceprevir, but its P4 group was clearly too large for the S4 pocket. Telaprevir could also be manually docked with good complementarity of its P1 sidechain and of its P2 group, which is a different proline analog from that of boceprevir (**Fig. 1C**), whereas its P4 group appeared slightly too large for the S4 pocket. The P3 group of coronavirus M^pro^ substrates face out into solution (*37*), as does the t-butyl group in the analogous position of boceprevir, telaprevir, and narlaprevir. Interestingly, the φ and Ψ angles of the proline ring in boceprevir precisely retraced the backbone atoms of 13b in the bound configuration (**Fig. 1D**). We thus hypothesized that boceprevir and, to a lesser extent, narlaprevir and telaprevir, may be able to inhibit SARSCoV2 M^pro^.

To address this hypothesis, we performed a preliminary rapid assessment using a live-cell assay of M^pro^ function. We co-expressed SARSCoV2 M^pro^ and a substrate protein comprising green fluorescent protein (GFP), a substrate site, and the membrane-targeting CAAX sequence (**Fig. 2A**). Cleavage at the substrate sequence liberates GFP, allowing quantitation of activity by immunoblotting. We could then assess the ability of drugs to inhibit M^pro^ activity in cells. In HEK293A cells, we found boceprevir and telaprevir to inhibit SARSCoV2 M^pro^ at micromolar concentrations (**Fig. 2B)**. These results provided evidence of the ability of HCV inhibitors with a P2 proline group to inhibit SARSCoV2 M^pro^.

**Figure 2.**
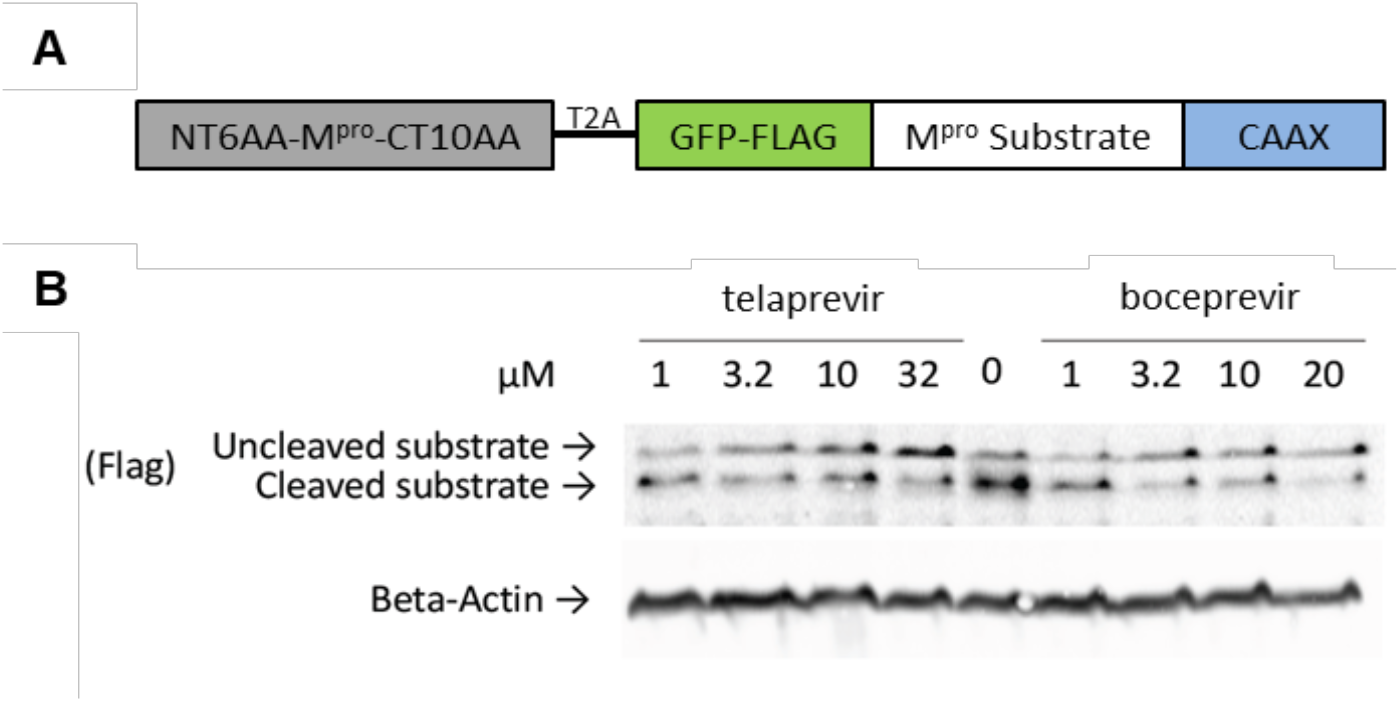
Preliminary evidence of HCV protease inhibitor activity on SARSCoV2 M^pro^. **(A)** Schematic of the protease reporter construct. An immature version of M^pro^ with reduced activity was used for this experiment to mimic multiple natural states of M^pro^. During the experiment, M^pro^ is expected to self-activate by removing the terminal extensions. **(B)** Immunoblot of lysates from HEK293A cells transiently transfected with the reporter construct shown in (A) and grown in the absence or presence of different concentrations of telaprevir or boceprevir shows suppression of substrate cleavage at micromolar drug concentrations.

We next performed a pilot test of the ability of HCV inhibitors with a P2 proline group to inhibit SARSCoV2 M^pro^ *in vitro* (**Fig. 3A**). As controls, we also tested GC-376 as a known broad-spectrum coronavirus M^pro^ inhibitor, ebselen and disulfiram as compounds recently reported to inhibit SARSCoV2 M^pro^ (*16*), and ritonavir as a protease inhibitor without any structural homology to coronavirus M^pro^ substrates. Indeed, we observed that boceprevir, narlaprevir, telaprevir, GC-376, disulfiram, and ebselen (at 150 μM concentration in air-equilibrated buffer) were able to inhibit SARSCoV2 M^pro^, whereas ritonavir showed no inhibitory effect (**Fig. 3A**). Incidentally, disulfiram and ebselen showed no inhibitory effect in the presence of DTT, whereas other compounds were unaffected by DTT. These results are consistent with proposals of disulfiram and ebselen forming a reducible bond with M^pro^ (*38*). As the interior of the cell is reducing, reduction of ebselen adducts could contribute to its relatively high EC_50_ of 4.7 μM for blocking SARSCoV2 replication in cells (*16*).

**Figure 3.**
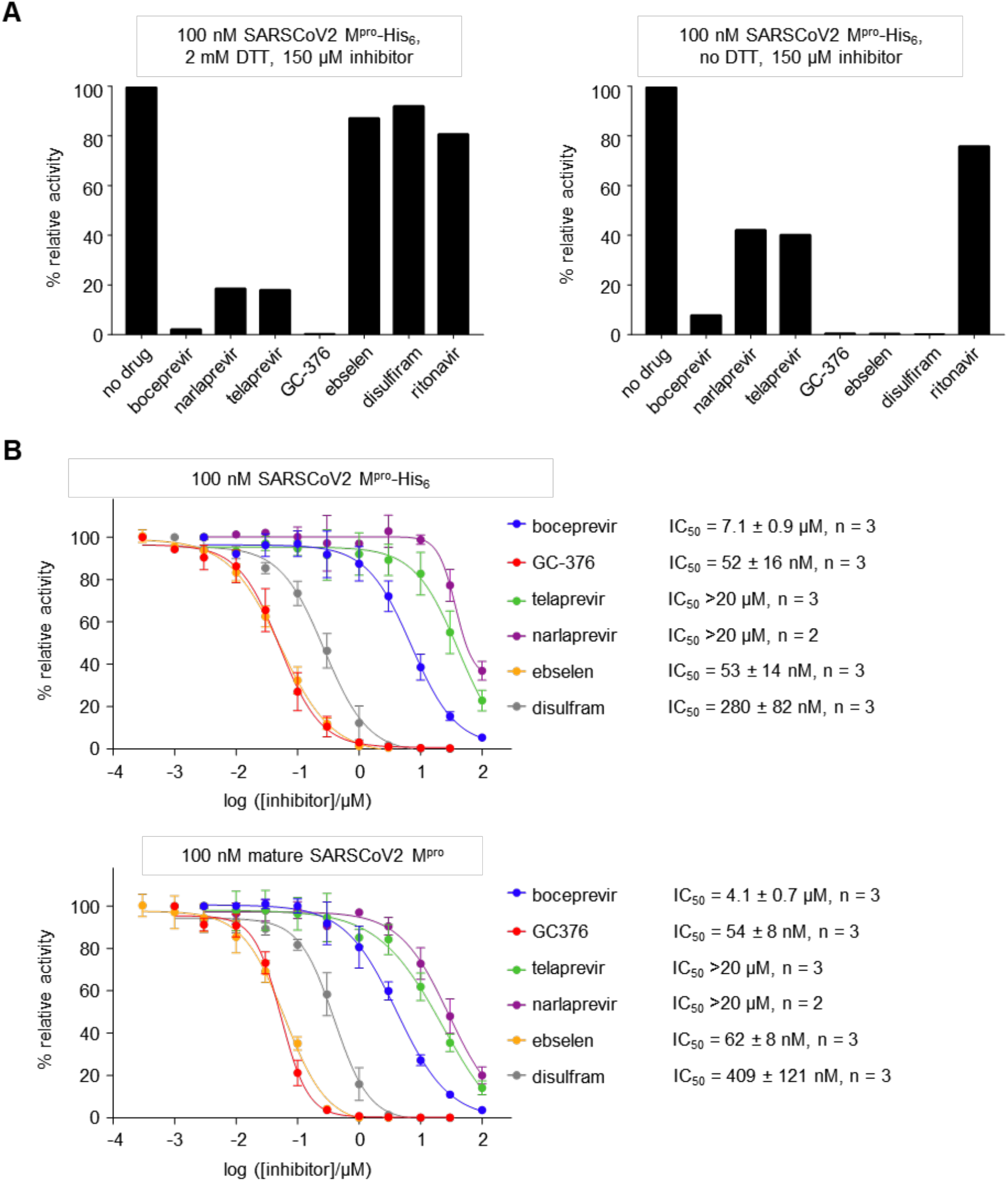
Inhibition of SARSCoV2 M^pro^ activity *in vitro*. **(A)** Left, relative M^pro^ activity in the presence of 2 mM DTT and 150 μM of drugs. GC-376 showed the most inhibition, followed by boceprevir, then telaprevir and narlaprevir. The HIV protease inhibitor ritonavir, ebselen, and disulfiram showed less than 50% inhibiting of enzyme activity, indicating IC_50_ > 150 μM in reducing conditions. Right, in the absence of DTT, ebselen and disulfiram showed efficient inhibition of M^pro^ activity. Enzymatic assays were carried out with 100 nM purified SARSCoV2 M^pro^ with an uncleaved C-terminal His_6_-tag, M^pro^-His_6_ **(B)** IC_50_ measurements by inhibitor titrations on 100 nM SARSCoV2 M^pro^-His_6_ (top) or 100 nM fully mature SARSCoV2 M^pro^ (bottom). For ebselen and disulfram, measurements were performed in the absence of DTT, while the assay buffer contained 2 mM DTT for all other drugs. Mean values of 2 to 3 independent experiments are shown. Error bars represent standard deviation.

We then performed full inhibition curves SARSCoV2 M^pro^ to obtain IC_50_ values for the above compounds. The initial pilot test above was performed with SARSCoV2 M^pro^ with a C-terminal His_6_-tag for rapid purification, but the presence of a C-terminal extension may have an inhibitory effect on protease activity (*39, 40*). We thus performed inhibition curves on both C-terminally extended and fully mature SARSCoV2 M^pro^ (**Fig. 3B**). We tested drugs in the presence of DTT, except for ebselen and disulfiram, for which we omitted DTT. Our results showed that the proline-containing HCV protease inhibitors boceprevir, telaprevir, and narlaprevir all could inhibit activity to some extent in both C-terminally extended and fully mature forms of SARSCoV2 M^pro^. Among these, boceprevir was the most potent, with an IC_50_ value of 4.1 μM on mature SARSCoV2 M^pro^ (**Fig. 3B, Table 1**).

**Table 1.**
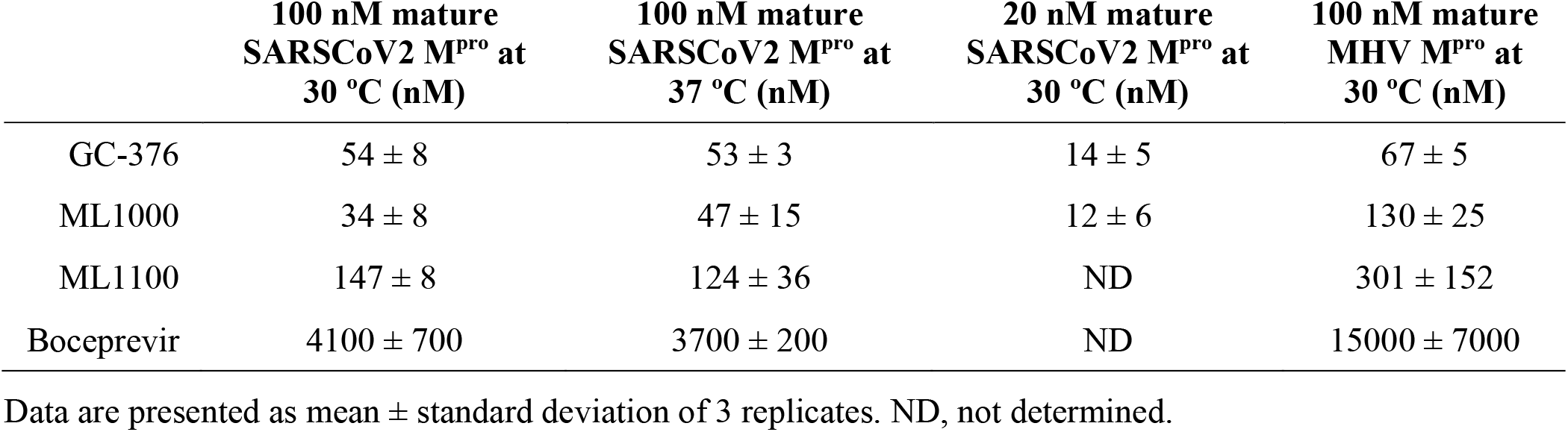
Measured 50% inhibitory constants (IC_50_ values) of ML1000 and ML1100 on SARSCoV2 M^pro^ *in vitro*

|  | 100 nM mature<br>SARSCoV2 M<sup>pro</sup> at<br>30 °C (nM) | 100 nM mature<br>SARSCoV2 M<sup>pro</sup> at<br>37 °C (nM) | 20 nM mature<br>SARSCoV2 M<sup>pro</sup> at<br>30 °C (nM) | 100 nM mature<br>MHV M<sup>pro</sup> at<br>30 °C (nM) |
| --- | --- | --- | --- | --- |
| GC-376 | 54 ± 8 | 53 ± 3 | 14 ± 5 | 67 ± 5 |
| ML1000 | 34 ± 8 | 47 ± 15 | 12 ± 6 | 130 ± 25 |
| ML1100 | 147 ± 8 | 124 ± 36 | ND | 301 ± 152 |
| Boceprevir | 4100 ± 700 | 3700 ± 200 | ND | 15000 ± 7000 |

Data are presented as mean ± standard deviation of 3 replicates. ND, not determined.

Having established that the P2 proline analogues in boceprevir, telaprevir, and narlaprevir were compatible with binding to the SARSCoV2 M^pro^ active site, we next sought to design next-generation coronavirus inhibitors that could combine the entropic stabilization conferred by these P2 rings with side groups optimized for coronavirus M^pro^ binding. A P1 Gln residue is strongly preferred by all coronavirus M^pro^ species (*14*). In fact, this preference is conserved with the related enterovirus 3C proteases such as human rhinovirus (HRV) protease and even with the more distantly related potyvirus proteases such as tobacco etch virus (TEV) protease (*41, 42*). Beginning with the HRV protease inhibitors AG7088 (rupintrivir) and its derivative AG7404 (*43, 44*), enterovirus and coronavirus protease inhibitors have incorporated a γ-lactam group to mimic the hydrogen bond acceptor function of Gln in the P1 position. Enzyme-inhibitor cocrystal structures have confirmed that the **γ**-lactam amide group is positioned similarly to the Gln amide group in natural structures (*37, 45*). We thus designed a novel inhibitor, ML1000, with the ketoamide warhead of boceprevir, the **γ**-lactamyl P1 group of rupintrivir, and the P2 through P4 moieties (including backbone and side chains) of boceprevir (**Fig. 4A**). We also designed a second inhibitor, ML1100, composed of the ketoamide warhead of boceprevir, the **γ**-lactamyl P1 group of rupintrivir, the P2 cyclic structure of telaprevir, and the P3 and P4 moieties of boceprevir (**Fig. 4A**). We retained the P3 and P4 structure of boceprevir as these groups did not demonstrate any clashes with the enzyme pocket in the manually docked boceprevir model, and as their hydrophobicity likely contributes to membrane permeability.

**Figure 4.**
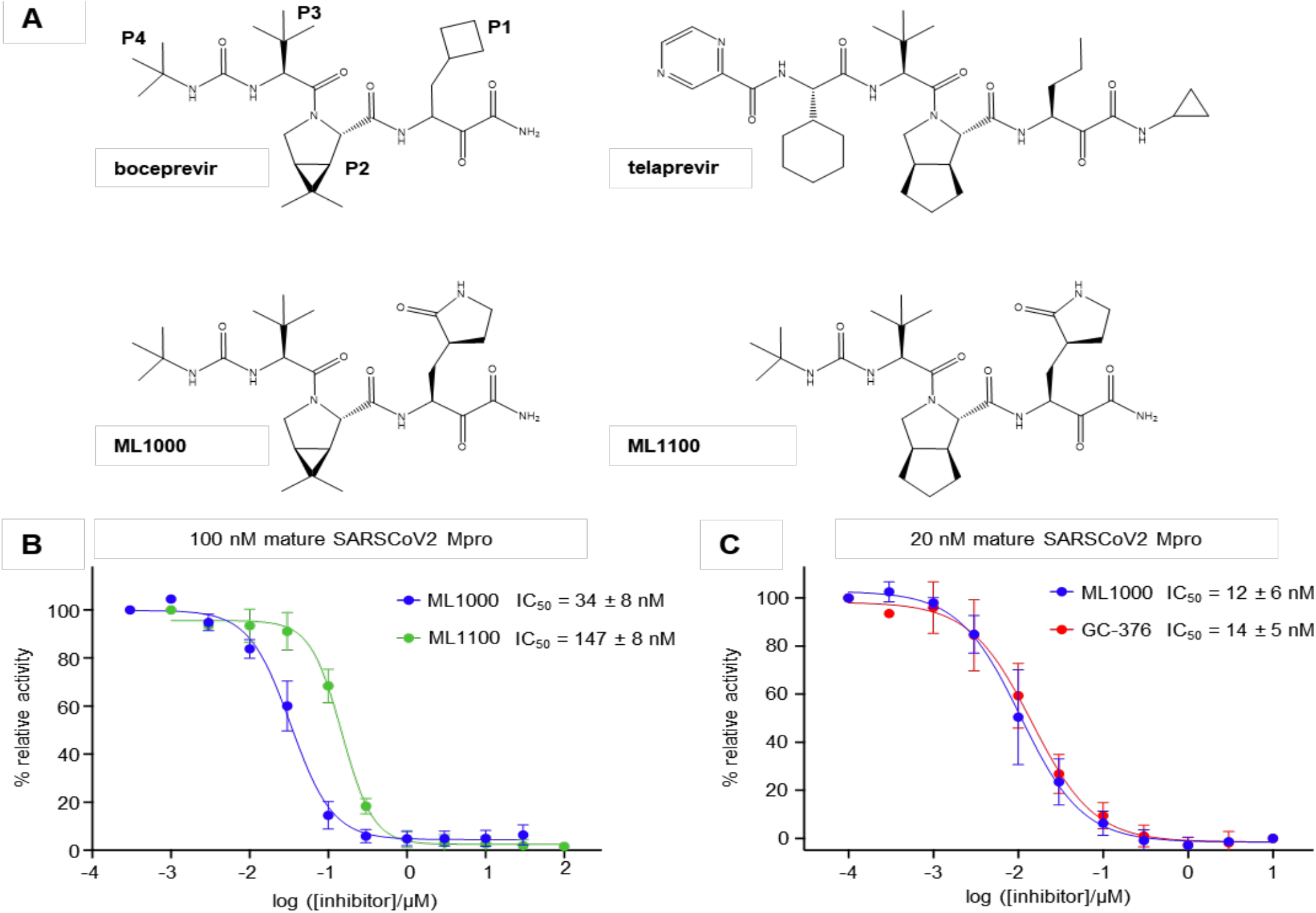
Design and *in vitro* potency of a novel coronavirus M^pro^ inhibitors. **(A)** Structures of boceprevir, telaprevir, ML1000, and ML1100. ML1000 is essentially boceprevir with a γ-lactamyl group in place of the P1 cyclobutanyl group. ML1100 replaces the bicyclic P2 proline analog of boceprevir with that of telaprevir. **(B)** IC_50_ measurements of ML1000 and ML1100 with 100 nM mature SARSCoV2 M^pro^. With enzyme concentration at 100 nM, the lowest possible IC_50_ that can be detected in theory is 50 nM. Thus, IC_50_ values were also measured for ML1000 and GC-376 with 20 nM enzyme. **(C)** IC_50_ values of ML1000 and GC-376 measured with 20 nM mature SARSCoV2 M^pro^ were 12 and 14 nM, respectively. Thus, ML1000 and GC-376 shows tight binding, even at 20 nM of M^pro^, and the IC_50_ values may still be limited by the enzyme concentration. Mean values of 3 independent experiments are shown. Error bars represent standard deviation.

ML1000 and ML1100 proved to be highly potent inhibitors of SARSCoV2 M^pro^. In the presence of 100 nM of enzyme, ML1000 and ML1100 produced IC_50_ values of 34 nM and 147 nM (**Fig. 4B, Table 1**), compared to 54 nM for GC-376 (**Fig. 3B, Table 1**). Against the M^pro^ enzyme from the distantly related coronavirus mouse hepatitis virus (MHV), ML1000 and ML1100 exhibited IC_50_ values of 130 and 301 nM, respectively, compared to 67 nM for GC-376 (**Table 1, Supporting Fig. 1**). These results demonstrate that ketoamide inhibitors with proline ring analogs at the P2 position can function as broad-spectrum coronavirus M^pro^ inhibitors with potency approaching that of aldehyde inhibitors.

We noted that the measured IC_50_ values of ML1000 and GC-376 for SARSCoV2 M^pro^ were within experimental error of the theoretical limit of half of the enzyme concentration in the assay (*46*), hindering the ability to discern differences in potency between them. We thus also measured IC_50_ values at a lower SARSCoV2 M^pro^ concentration of 20 nM **(Fig. 4C)**. In these conditions, ML1000 and GC-376 still demonstrated comparable IC_50_ values of 12 and 14 nM, respectively, below the concentration of M^pro^ (i.e. the tight binding limit). Physiological temperature, compared to the 30 °C at which these assays are usually performed, could potentially increase structural flexibility and alter drug affinity of M^pro^. However, we found no significant difference in potency between 37 °C and 30 °C for each inhibitor (**Table 1, Supporting Fig. 2**). These results indicate that ML1000 has high potency *in vitro* and represents the tightest-binding non-aldehyde SARSCoV2 M^pro^ inhibitor discovered so far.

Next, we tested the efficacy of the new inhibitors against SARSCoV2 M^pro^ in human Huh7 cells by measuring the extent of cleavage of coexpressed substrate (**Fig. 5**). Here, we observed the rank order of effectivenss to be GC-376 > ML1000 > ML1100 > boceprevir (**Fig. 5**). It should be emphasized that this assay does not directly measure the fraction of active M^pro^. If a small fraction of active M^pro^ is sufficient to result in substrate cleavage, this would effectively increase the measured concentration needed to suppress protease activity. However, the assay does provide an assessment of the relative ability of a set of drugs to cross the cellular membrane and inhibit M^pro^ within living cells.

**Figure 5.**
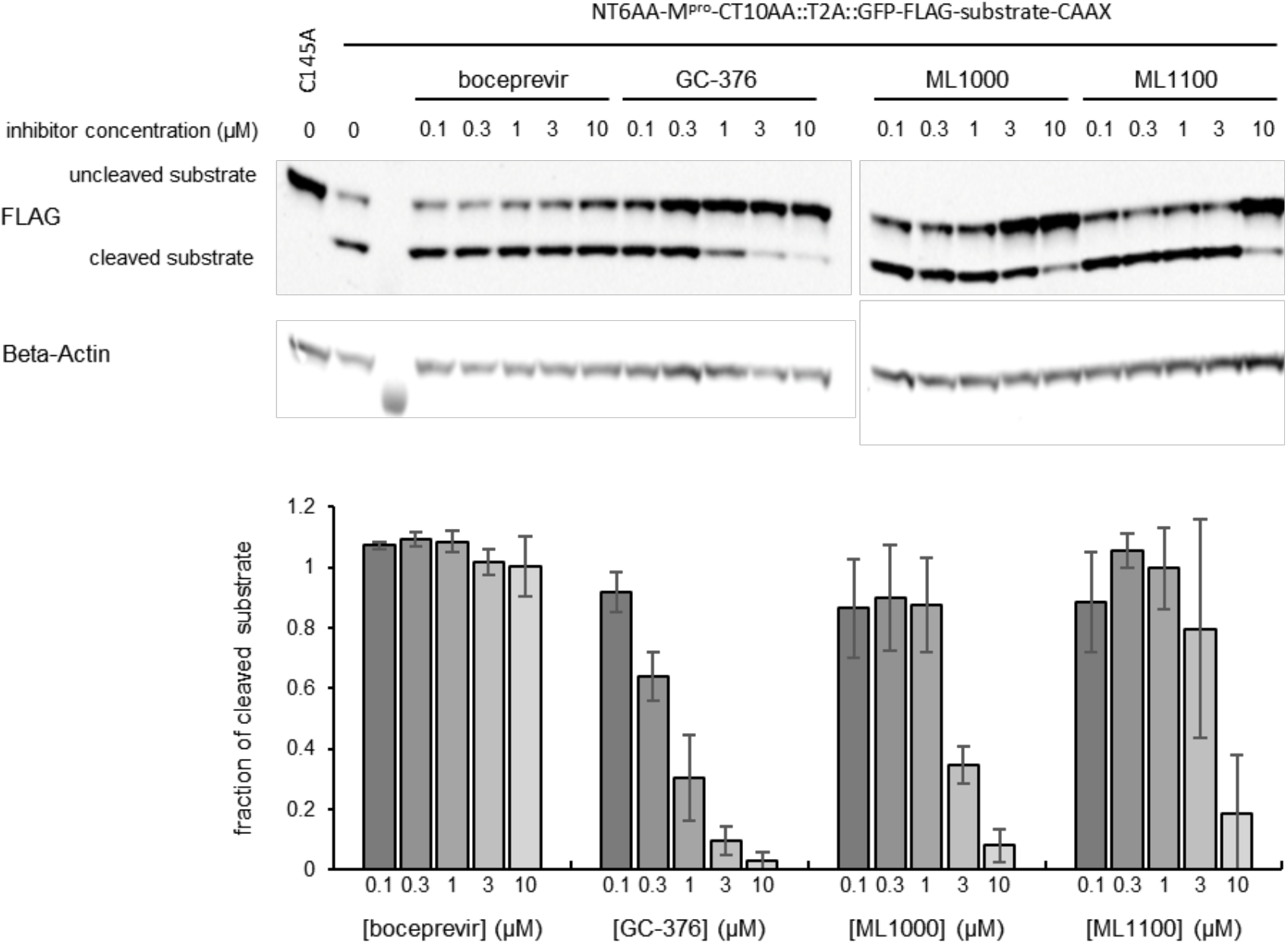
Potency of ML1000 and ML1100 in inhibiting M^pro^ activity in Huh7 cells. Above, representative immunoblot of lysates of Huh7 cells transduced with lentivirus encoding the construct shown in **Fig. 2A**. Immunoblots were probed with antibodies against FLAG to detect the substrate protein and beta-actin as a control. Below, the fraction of cleaved substrate relative to the total substrate abundance was quantified at different inhibitor concentrations and normalized to the cleavage ratio in absence of inhibitor. Mean values of 3 independent experiments are shown. Error bars represent standard deviation. The relative potency of the drugs in this assay is GC-376 > ML1000 > ML1100 > boceprevir.

Finally, we tested the ability of ML1000 and ML1100 to inhibit SARSCoV2 virus replication in Caco-2 human intestinal cells. For comparison, we tested boceprevir and GC-376. Boceprevir inhibited viral replication with an EC_50_ of 0.2 μM, while GC-376 was even more potent, with a remarkably low EC_50_ of 0.1 nM (**Table 2, Fig. 6**). ML1000 and ML1100 inhibited SARSCoV2 replication with EC_50_ values of 0.1 and 0.2 μM, respectively (**Table 2, Fig. 6**). While the enhancement in anti-viral efficacy of ML1000 relative to boceprevir is not as large as the enhancement in enzyme inhibition observed *in vitro* (**Table 1**), the EC_50_ value of ML1000 is nevertheless the lowest reported to date for any non-aldehyde inhibitors of SARSCoV2 M^pro^. In addition, ML1000 and ML1100 exhibited no cytotoxicity at the highest dose tested of 100 μM in Caco-2 cells and Huh-7 cells (**Table 2, Supporting Fig. 3**), resulting in selectivity indices of >1000 and >500 respectively. These results confirm that ML1000 is a potent and selective ketoamide inhibitor of SARSCoV2 replication.

**Table 2.** Antiviral activity of ML1000 and ML1100 on SARSCoV2 replication in Caco-2 cells

|  | EC<sub>50</sub> (μM) | CC<sub>50</sub> (μM) | Selectivity index |
| --- | --- | --- | --- |
| GC-376 | 0.0001 | >100 | >10<sup>6</sup> |
| ML1000 | 0.1 | >100 | >1000 |
| ML1100 | 0.2 | >100 | >500 |
| Boceprevir | 0.2 | >100 | >500 |

EC<sub>50</sub>: 50% effective antiviral concentration
CC<sub>50</sub>: 50% cytotoxic concentration of compound without virus added
Selectivity index = CC<sub>50</sub>/EC<sub>50</sub>

**Figure 6.**
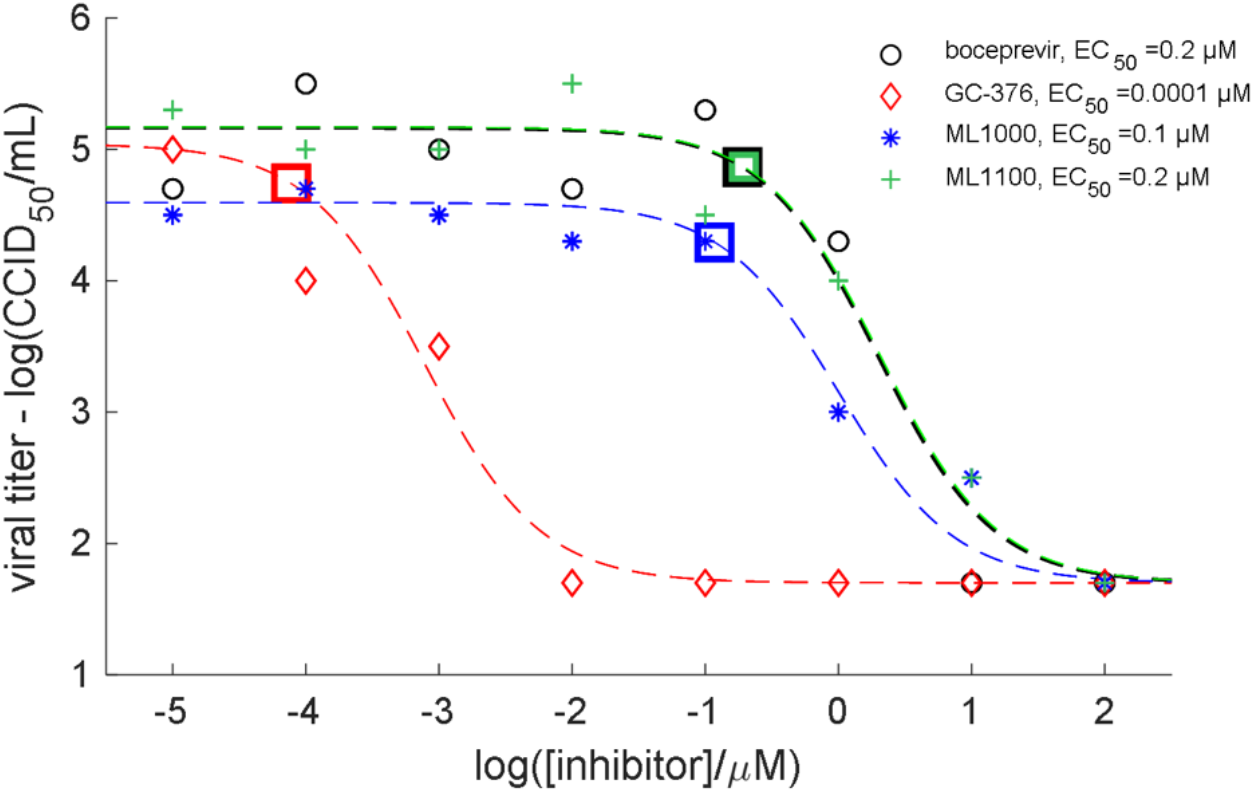
Inhibition of SARSCoV2 replication in Caco-2 cells. The viral titer was quantified after incubation with different concentrations of inhibitor. For each condition, three technical replicates were combined for a single measurement. The data were plotted on a log-log graph and fit to a logistic function. EC_50_ values were calculated as the inhibitor concentration needed to reduce the viral titer by one log2 unit relative to the upper baseline, and the corresponding points on the curves are indicated with squares on each fitted curve.

## DISCUSSION

There is strong interest in discovering non-aldehyde inhibitors of SARSCoV2 M^pro^ with the potential for oral administration. In this study, we developed ML1000 as a ketoamide inhibitor that is more potent against SARSCoV2 replication than previously described ketoamide or Michael acceptor compounds (*9*). Although its safety profile remains to be determined, ML1000 was designed to minimize toxicity by using only chemical motifs from drugs with established safety records in humans. Likewise, ML1000 was designed with the potential for oral dosing in mind. ML1000 is essentially the approved drug boceprevir with its P2 group altered to that of AG-7404. Of note, both boceprevir and AG-7404 have been proven to have high oral bioavailability and to be safe in humans (*44, 47*). In particular, boceprevir is taken 3 times daily for over 300 consecutive days on an outpatient basis (*47*).

Our findings that GC376 potently inhibits SARSCoV2 replication in Caco-2 cells adds to other recent data supporting its potential utility for treating COVID-19 (*20, 48, 49*). However, GC376 may be better suited for intravenous administration than oral dosing, given its short *in vivo* half-life and low oral bioavailability (*12, 27*). In addition, the presence of off-target effects, evidenced by suppression of tooth eruption in GC376-treated cats (*13*), suggests the need to develop additional classes of protease inhibitors.

In the course of our study, three other groups reported that boceprevir can inhibit SARSCoV2 M^pro^ at low micromolar concentrations *in vitro* (*48–50*). Two of these studies included crystal structures of SARSCoV2 M^pro^ complexed with boceprevir (PDB 6WNP and 7C6S) that validate our manually docked model (*49, 50*). Alignment of these structures to that of 13b complexed with SARSCoV2 M^pro^ indicates that the proline ring of boceprevir can indeed mimic a turn taken by the backbone of linear protease inhibitors at the P2 position (**Supporting Fig 4**), confirming our original hypothesis (**Fig. 1**). Two of the studies also demonstrated that boceprevir suppresses SARSCoV2 viral replication in monkey cells with EC_50_ values between 2 and 16 μM (*48, 50*).

Our work demonstrates the unexpected favorability of incorporating proline-like rings into coronavirus protease inhibitors, specifically at the P2-analogous position. This finding is unexpected because proline is never found at this position in coronavirus M^pro^ substrates (*14*). One explanation could be that proline itself naturally lacks the bulk to fully occupy the deep S2 pocket, which prefers long aliphatic side chains such as leucine and phenylalanine. Thus, the proline analogs of boceprevir and telaprevir, which have a bicyclic structure, not only lock the backbone of the inhibitor into its bound conformation, but through their large bulk also fill the S2 pocket.

The ability of ML1000 and its structural variant ML1100 to bind to SARSCoV2 M^pro^ more effectively than larger compounds with the same ketoamide warhead (*17*) suggests that introduction of a cyclic structure can be a chemically parsimonious method to improve molecular recognition. The pre-rigidification of an inhibitor using a cyclic structure may provide enhanced affinity compared to a non-cyclic structure of the same size through prepayment of the entropic penalty of binding, and therefore can be an efficient strategy for potency improvement.

One parameter for potential further improvement of ML1000 is cell permeability. The conversion of the cyclobutyl group in the P1 position of boceprevir to the γ-lactamyl group in ML1000 adds an additional hydrogen bond acceptor and donor, which would be expected to decrease cell permeability. While ML1000 is ∼340-fold more potent on enzyme inhibition than boceprevir *in vitro*, it is only ∼2-fold more effective on viral suppression in Caco-2 cells. We speculate that improving the permeability of ML1000 would further enhance antiviral efficacy in cells.

In summary, ML1000 can be considered an example of a new class of coronavirus M^pro^ inhibitors that feature a cyclic P2 group, high affinity, and solubility. Taken together, our results suggest that ML1000 is a promising scaffold for the development of anti-coronavirus drugs, and additional structures incorporating proline ring analogues can be explored as coronavirus M^pro^ inhibitors.

## METHODS

### Inhibitors

ML1000 and ML1100 were synthesized under fee-for-service agreements by ACME Bioscience (Palo Alto, CA, USA), and Chempartner (Shanghai, China), respectively. All other inhibitors were readily available: boceprevir (Cayman Chemical, ≥ 98%), narlaprevir (AdooQ, ≥ 98%), telaprevir (AdooQ Bioscience, ≥ 98%), GC-376 (AOBIOUS, ≥ 98%), ebselen (Cayman Chemical, ≥ 99%), disulfiram (LKT Laboratories, ≥ 98%), ritonavir (Santa Cruz Biotechnology, ≥ 98%).

### Cell-based M^pro^ Activity Assay in HEK293A and Huh7

#### Cell culture and transfection

HEK293A and HEK293FT cells were cultured at 37 °C in 5% CO_2_ in Dulbecco’s Modified Eagle’s Medium (DMEM, Gibco) supplemented with 10% FBS and 100 U/mL penicillin and 100 μg/mL streptomycin. Huh7 cells were cultured at 37 °C in 5% CO_2_ in Roswell Park Memorial Institute 1640 medium (RPMI 1640, Life Technoloiges) supplemented with 10% FBS and 100 U/mL penicillin and 100 μg/mL streptomycin.

#### M^pro^ activity assay in HEK293A

Cells were transfected with a pcDNA3.1/Puro-CAG plasmid containing the construct shown in **Fig. 2A** using Lipofectamine 3000 (Life Technologies) in Opti-MEM (Life Technologies) according to the manufacturer’s protocol. Telaprevir and boceprevir were added 2 h post-transfection. 24 h post-transfection, cells were washed twice with PBS, then lysed with 50 μL hot LDS lysis buffer (50% 4x LDS Sample Buffer (NuPAGE, Life Technologies), 10% 2-mercaptoethanol), and DNA was sheared by sonication. After heating at 80-90 °C for 1-2 minutes, cell lysates were loaded onto 4%-12% or 12% Bis-Tris gels (NuPAGE, Life Technologies) along with a Precision Plus protein dual-color standard (Bio-Rad). Protein bands were transferred to PVDF membranes using a Trans-Blot Turbo Transfer System (Bio-Rad) and blocked with SuperBlock T20 (TBS) Blocking Buffer (Thermo Scientific). Membranes were probed with primary antibodies in SuperBlock T20 (TBS) Blocking Buffer and fluorophore-conjugated secondary antibodies in SuperBlock (TBS) Blocking Buffer (Thermo Scientific), with 3 washes in SuperBlock T20 (TBS) Blocking Buffer after each step. Membranes were imaged using an Odyssey imaging system (LI-COR). Western blots were quantified using ImageJ. The following primary antibodies were used for immunoblotting at the indicated dilutions: mouse monoclonal anti-FLAG (Sigma-Aldrich, F1804), 1:2000; rabbit polyclonal anti-beta Actin (Abcam, ab8227), 1:5000. Secondary antibodies were LI-COR 680RD goat-anti-rabbit and 800CW goat-anti-mouse, used at 1:5000 dilution each.

#### M^pro^ activity assay in Huh7

Lentiviruses were produced by transfecting HEK293FT cells with pLL3.7 plasmids containing the construct shown in **Fig. 2A** or a version with a catalytically dead M^pro^ C145A mutation and two helper plasmids (psPAX2, MD2G). Transfection was carried out with CalPhosTM Mammalian Transfection Kit (Takara Bio) at a ratio of pLL3.7:psPAX2:MD2G = 20:15:6. The virus-containing supernatant was harvested 48 or 72 hours post transfection. Cell debris was removed with a 5 min 2000 g centrifugation, followed by a 0.45 μm filtration (Millipore). Virus titer was determined using Lenti-XTM GoStixTM Plus (Takara Bio), and Huh7 cells were infected at a MOI of 5. Drugs were added 24 hours post infection and cells were lysed and prepared for western blotting, as described above, 4 days post infection.

### M^pro^ Expression and Purification

Modified versions of previous protocols based on HRV protease (*40*) or SUMO protease (*51*) processing of M^pro^ fusion proteins were used to obtain purified M^pro^ with either native or extended termini after expression in E. coli. Specifically, M^pro^ variants were cloned either as a GST-M^pro^-His_6_ or His_6_-SUMO-M^pro^ fusion into a pET vector (Addgene plasmid #29666) using synthetic gene blocks for the M^pro^ portion of the SARS-CoV2 polyprotein (pp1ab residue 3264-3569) or the MHV1 polyprotein (pp1ab 3314-3624). The constructs are listed in **Supporting Fig. 5**. In general, the His_6_-SUMO-M^pro^ fusions produced higher yields of soluble protein compared to the GST-M^pro^-His_6_ constructs. This is likely due to toxicity and growth retardation associated with M^pro^ activation upon autocatalytic removal of the GST-tag during expression of the GST-M^pro^-His_6_ fusion. In contrast, the His_6_-SUMO-M^pro^ fusion is produced in full-length and only becomes fully active after SUMO-tag removal during subsequent purification steps.

For SARSCoV2 M^pro^ variants, the plasmids were transformed into T7 Express lysY/I^q^ Competent E. coli (NEB). All cultures were grown in 2×YT media. Overnight cultures were used to inoculate larger cultures that were grown at 37 °C to OD_600_ ∼ 0.8 before induction with 0.5 mM IPTG. After induction and 4-5 h growth at 24 °C, the cells were harvested, and the pellets were frozen. Chemical lysis of the resuspended pellets was performed in BPER (Thermo Fisher) supplemented with 20 U/mL Pierce universal nuclease (ThermoFisher Scientific), and the supernatant cleared by 15 min of centrifugation at 15000 g. The soluble fraction was batch absorbed onto INDIGO-Ni resin (Cube Biotech) in a buffer with imidazole and NaCl added to 10 mM and 200 mM, respectively. The resin was loaded onto gravity flow columns and washed with 20 column volumes of wash buffer containing 50 mM Tris (pH 8), 25 mM Imidazole, and 300 mM NaCl. High purity protein was eluted in a buffer of 50 mM Tris (pH 8), 250 mM Imidazole, and 300 mM NaCl. Eluted fractions with high protein content were pooled and buffer exchanged into HRV protease cleavage buffer (50 mM Tris (pH 7.3), 150 mM NaCl, 1 mM DTT) or SUMO protease cleavage buffer (50 mM Tris (pH 8), 150 mM NaCl, 1 mM DTT). The proteins were cleaved by incubation with 1.5 % (w/w) His-tagged HRV-3C protease (Millipore Sigma SAE0045) or 15 U/mg His-tagged SUMO protease (Millipore Sigma SAE0067) by overnight incubation at 4 °C. The fully processed M^pro^ variants were then purified using reverse affinity chromatography to remove the His-tagged HRV and SUMO proteases and the cleaved His_6_-tagged fusion-domains/peptides. Purity of the samples was checked on SDS-PAGE, and protein concentrations were determined based on A_280_ and predicted extinction coefficients.

For MHV M^pro^, expression from construct IV (**Supporting Fig. 5**) did not result in acceptable yields of M^pro^-His_6_. Furthermore, we found that the expression of construct V and VI could be dramatically improved by adding 10 mM DTT to the lysis, wash, and elution buffers described above. All other steps and conditions were unchanged from the protocol described above for SARSCoV2 M^pro^.

### In-Vitro M^pro^ Inhibition Assays

The proteolytic activity of purified M^pro^ was primarily measured using a fluorogenic peptide, Covidyte IF670 (AAT Bioquest), that includes a far-red fluorophore iFluor™ 670 and a quencher Tide Quencher™ 5, TQ5. Upon cleavage of the iFluor-VNSTLQ/SGLRK(TQ5)M peptide by M^pro^, energy transfer to the quencher decreases and the fluorescence intensity of iFluor increases. A well-plate reader (Tecan, Safire 2) was used to monitor the fluorescence with excitation at 640/20 nm and emission at 680/20 nm. All measurements were performed as bottom-reads from lid-covered black 96 well plates with clear bottoms. The in-vitro M^pro^ activity was measured in an assay buffer consisting of 50 mM Tris (pH 7.3), 50 mM NaCl, 1 mM EDTA, 2 mM DTT, and 1-2% DMSO. In specific experiment DTT was omitted from the buffer.

For the majority of IC_50_ measurements, 100 nM M^pro^ was preincubated with varying concentrations of inhibitor for 30 min at 30 °C. Next, the substrate was added to a final concentration of 2 μM and a final volume of 90 μL. The well-plate was immediately inserted into the plate reader, that had been preheated to 30 °C, and the fluorescence intensity was monitored at 30 s intervals for 45 min. For selected experiments, the measurements were performed at 37 °C instead of 30 °C. For GC-376 and ML1000, IC_50_ values were also quantified at a lower M^pro^ concentration of 20 nM in attempt to escape the tight binding regime (**Supporting Fig. 6C**). For these experiments, a substrate with faster reaction rate, Covidyte TF670 (AAT Bioquest), was used to counteract the reduced M^pro^ activity and increase the sensitivity of the assay. This substrate consists of a far-red fluorophore Tide Fluor™ 5, TF5, and the TQ5 quencher linked by a peptide substrate, TF5-KTSAVLQ/SGFRKME(TQ5)M.

The rate of substrate cleavage was extracted by linear fitting of the fluorescence signal increase as a function of time (**Supporting Fig. 6**). The IC_50_ curves were evaluated using the following equation in Prism (GraphPad):

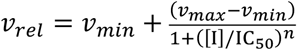

Here *v*_rel_ is the experimentally measured rate of substrate cleavage normalized to the rate of substrate cleavage in the absence of inhibitor, *I*. The maximal and minimal rate of substrate cleavage in each experiment, *v*_max_ and *v*_min,_ respectively, the Hill coefficient, *n*, and IC_50_ are all fitting parameters in the non-linear fitting routine.

### Cytotoxicity assay in Huh7 cells

24 h prior to the drug treatment, 10000 Huh7 cells were seeded in 100 μl culture medium (DMEM with 10% FBS, 2mM L-glutamine, penicillin-streptomycin) in a 96-well white microplate (Greiner Bio-One, Austria) pre-coated with poly-D-lysine (10μg/ml, Sigma). The next day, the culture medium was replaced with assay medium (DMEM with 5% FBS, 2mM L-glutamine, penicillin-streptomycin, 1 % DMSO) containing inhibitors at the desired concentration (1 −100 μM). Staurosporine, a non-selective protein kinase inhibitor known to induce apoptosis, was used as a positive control (0.01 μM - 1 μM). After 48 h, cell viability was determined using the CellTiter-Glo 2.0 kit (Promega, USA) according to the instructions of the manufacturer. The bioluminescence signal was measured on a multi-mode microplate reader (Tecan, Safire 2).

### Antiviral and cytotoxicity assays in Caco-2 cells

The tested M^pro^ inhibitors were serially diluted from 10 mM DMSO stocks using eight log_10_ dilutions in test medium (MEM supplemented with 2% FBS and 50 μg/mL gentamicin) yielding a concentration range of 10 pM – 100 μM. Each dilution was added to 5 wells of a 96-well plate with 80-100% confluent Caco-2 cells. Three wells of each dilution were infected with virus, and two wells remained uninfected as toxicity controls. Six wells were infected and untreated as virus controls, and six wells were uninfected and untreated as cell controls. SARSCoV2 (USA_WA1/2020 strain passaged twice in Vero 76 cells in MEM supplemented with 2% fetal bovine serum and 50 μg/ml gentamicin to prepare a working stock) was prepared at a multiplicity of infection (MOI) that would yield measurable virus titers within 72 hours. Plates were incubated at 37±2 °C and 5% CO_2_.

For virus yield reduction assays, the supernatant fluid from each condition was collected on day 3 post-infection (3 wells pooled) and tested for virus titer using an endpoint dilution in Vero 76 cells. The virus titer was determined by visual observation of cells under a light microscope on day 5 post-infection. Viral titers, *VT*, were quantified on a logarithm scale in the form of CCID_50_/mL using the Reed Muench equation. The data was fitted to the following equation:

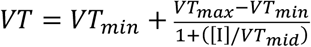

The 50% effective concentration, EC_50_, was defined as the concentration of inhibitor were *VT* reaches *VT*_*max*_ −log2 and extracted from the fitted curves.

Drug cytotoxicity was also assayed on day 3 in a neutral red viability assay. Plates were stained with dye for 2 hours (±15 minutes). Supernatant dye was removed, wells rinsed with PBS, and the incorporated dye extracted in 50:50 Sørensen citrate buffer/ethanol for >30 minutes before measuring the optical density at 540 nm, OD_540_. OD_540_ was used to calculate the relative viability to non-exposed cell controls.

### Significant hazards

No unexpected or unusually high safety hazards were encountered.

## ACKNOWLEDGEMENTS

The pET His6 TEV LIC cloning vector (2B-T) was a gift from Scott Gradia (Addgene plasmid # 29666; http://n2t.net/addgene:29666; RRID:Addgene_29666). The project is supported by a ChEM-H accelerator grant for COVID-19 from Stanford ChEM-H, and a Fast Grant for COVID-19 from Emergent Ventures at the Mercatus Center at George Mason University. M.W. is funded by grant NNF18OC0031816 from the Novo Nordisk Foundation and the Stanford Bio-X Program. X.Z. is funded by a Stanford Bio-X SIGF Graduate Student Fellowship.

---

**Supporting Figure 1.**
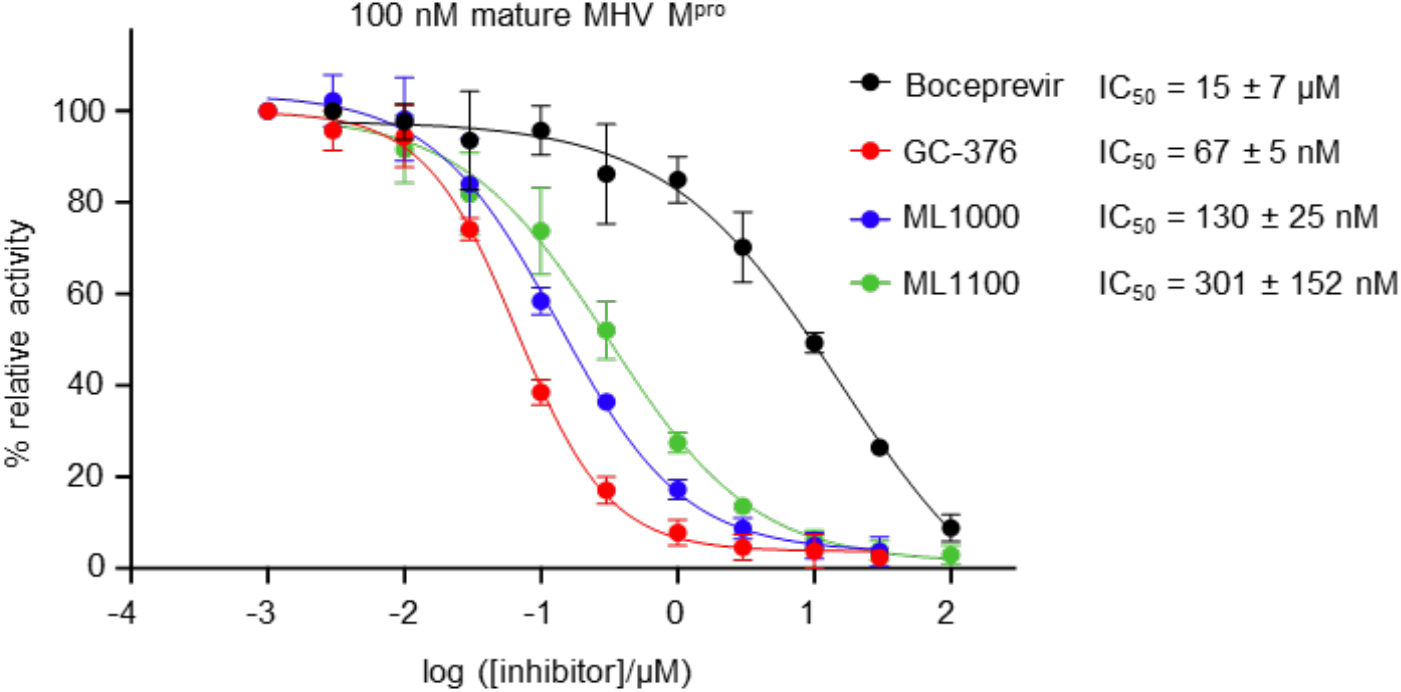
IC_50_ measurements for boceprevir, GC-376, ML1000, and ML1100 with 100 nM mature MHV M^pro^ at 30°C. Three independent experiments were performed. Error bars represent standard deviation.

**Supporting Figure 2.**
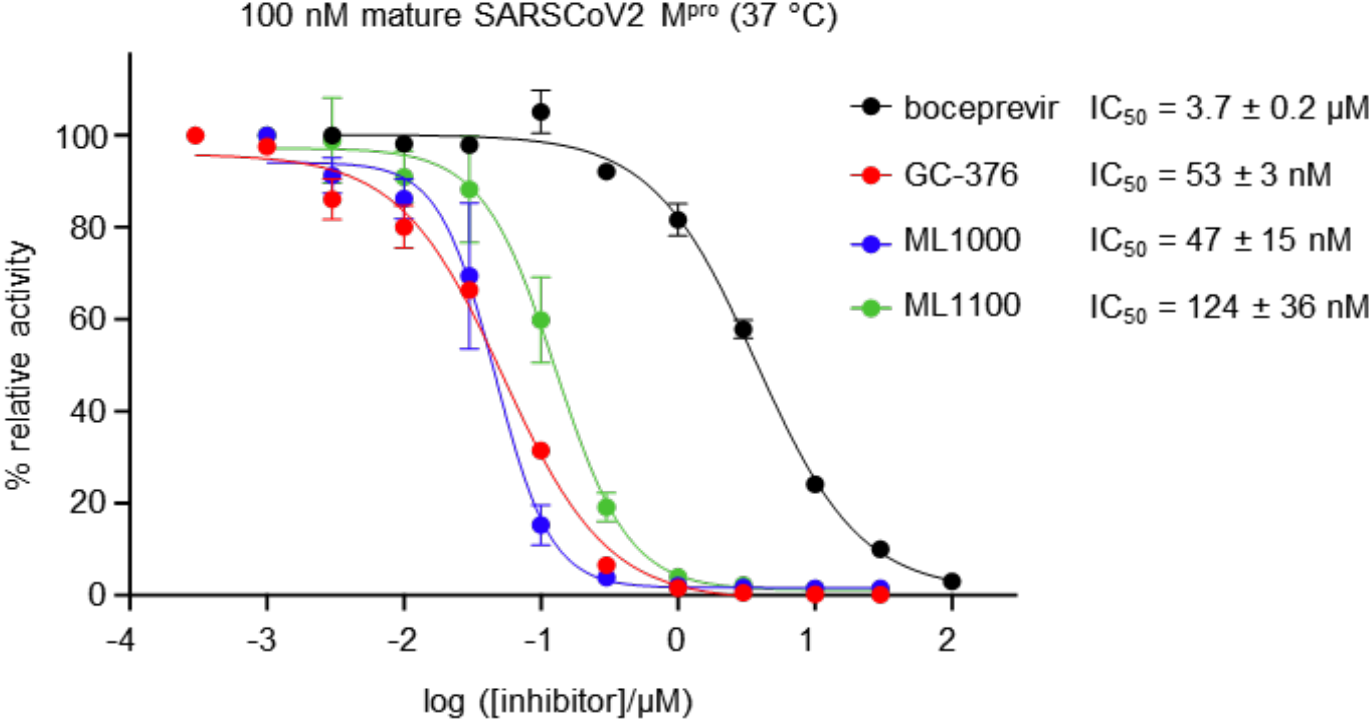
IC_50_ measurements for boceprevir, GC-376, ML1000, and ML1100 with 100 nM mature SARSCoV2 M^pro^ at 37°C. Three independent experiments were performed. Error bars represent standard deviation.

**Supporting Figure 3.**
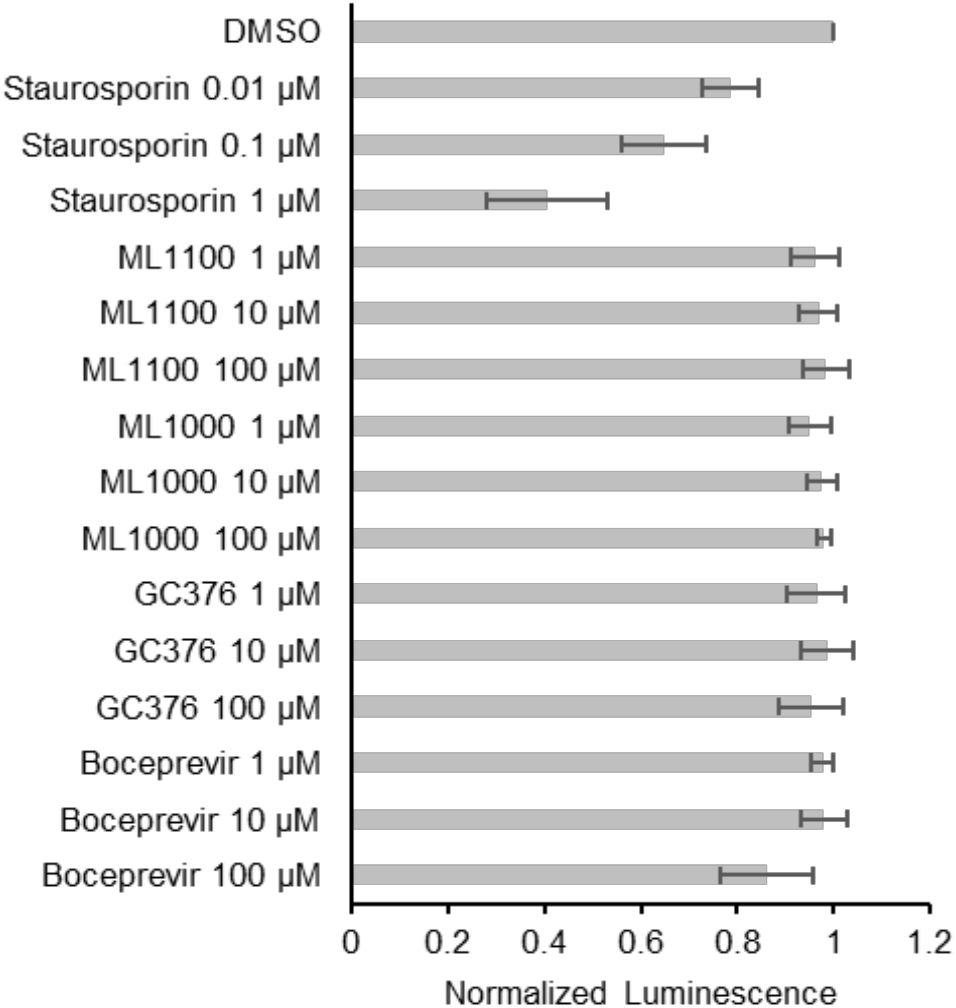
Testing of protease inhibitor compounds on Huh-7 cells in a viability assay. Luminescence signals from CellTiter-Glo 2.0 reagent were normalized to DMSO. Staurosporine was used as a control cytotoxic compound. Three independent experiments were performed. Error bars represent standard error of the mean.

**Supporting Figure 4.**
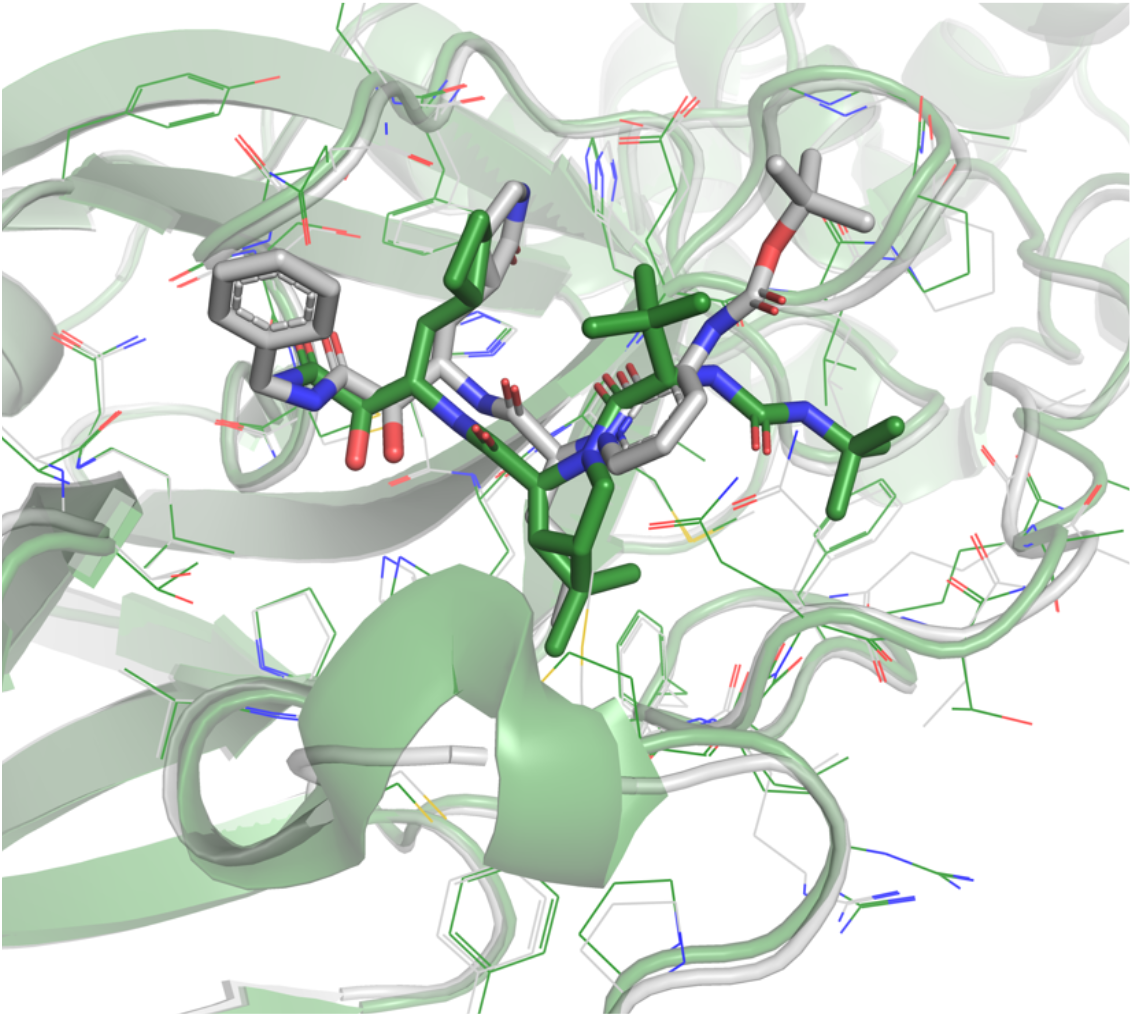
Alignment of PDB structures 6Y2F (grey) and 6WNP (green) shows that the proline-like kink at P2 is indeed a common structural motifs of the inhibitors 13b (grey) and BPV (green).

**Supporting Figure 5.**
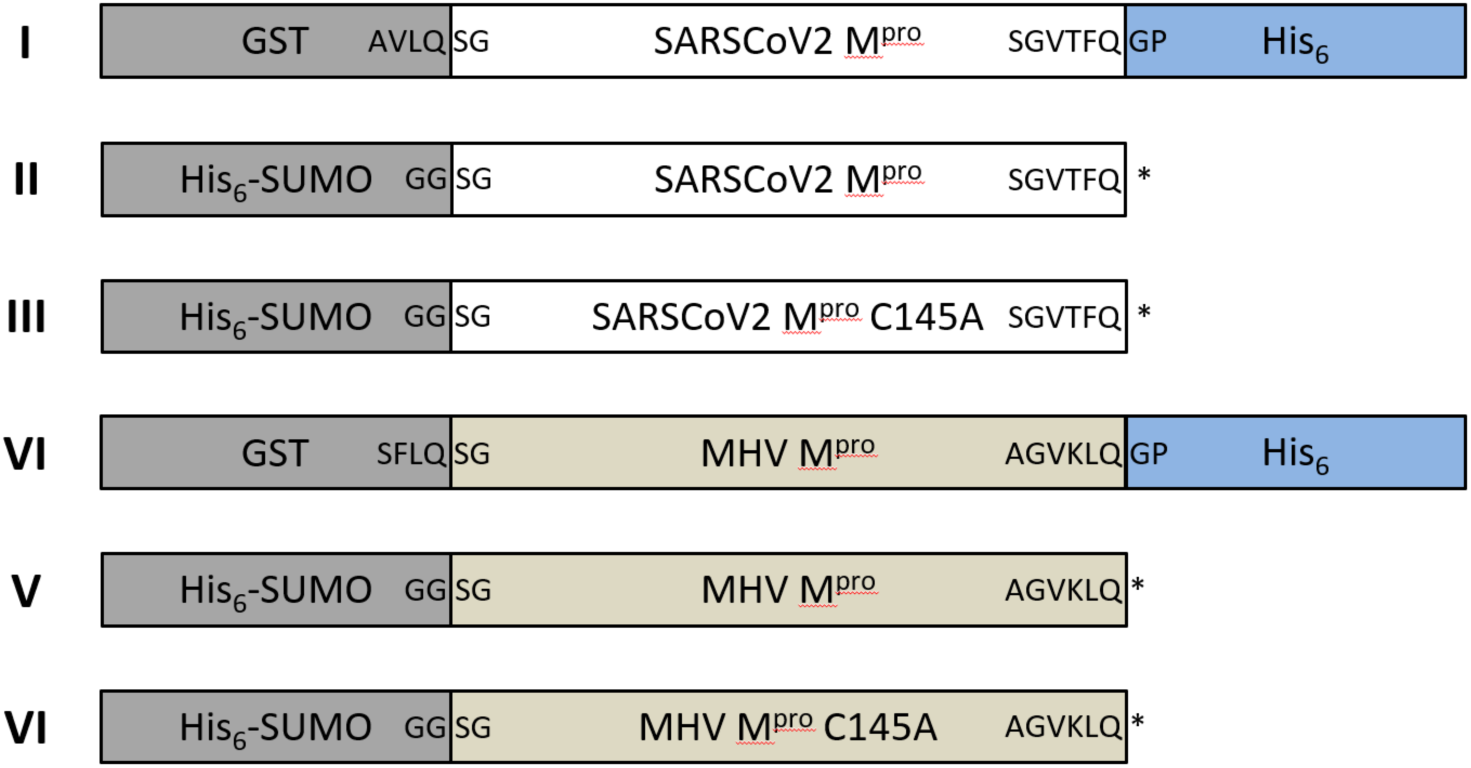
M^pro^ constructs for expression in E. coli. The fusion constructs are listed with the cleavable amino acid sequences highlighted at the domain boundaries. For construct I and IV, the N-terminal GST is autocatalytically removed during expression, while the C-terminal His_6_-tag was removed during purification. Selected experiments (**Fig. 3**) were performed with construct I without removal of the His_6_-tag. After HRV and SUMO protease cleavage, respectively, fully mature SARSCoV2 M^pro^ from either construct I or II performed similarly in all assays, but protein expressed from II was used for data on mature SARSCoV2 M^pro^ from reported here.

**Supporting Figure 6.**
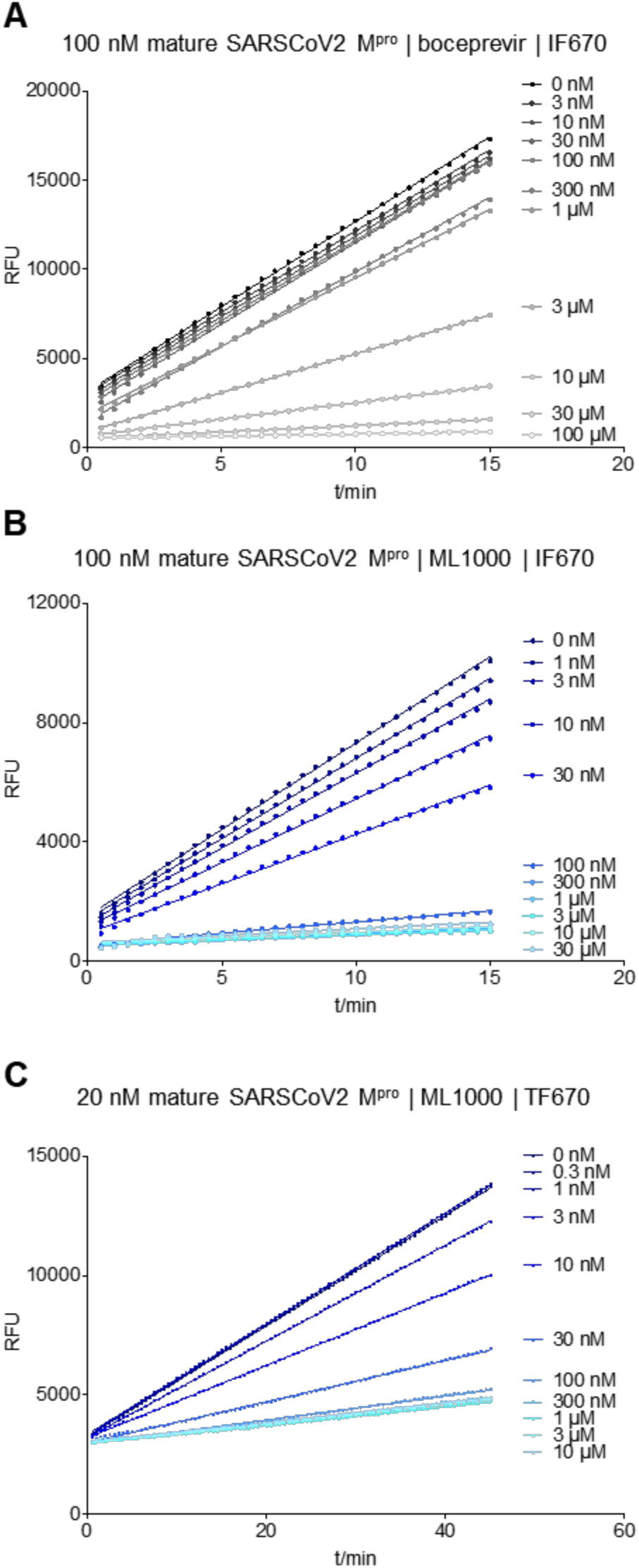
Representative datasets for *in vitro* cleavage of fluorogenic substrates by mature SARSCoV2 M^pro^. The time-resolved fluorescence intensity increases due the proteolytic cleavage of Covidyte IF670 by 100 nM mature SARSCoV2 M^pro^ was monitored as a function of inhibitor concentration: **(A)** boceprevir and **(B)** ML1000. **(C)** For ML1000, the assay was also performed at lower enzyme concentrations and with a different substrate (Covidyte TF670) to allow for better quantification of tight inhibition. In all cases, the rate of substrate cleavage was extracted as the slope of linear fits. Catalytically dead M^pro^ C145A **(Supporting Fig. 5**) was used to establish the baseline the assays.

